# Simulations suggest walking with reduced propulsive force would not mitigate the energetic consequences of lower tendon stiffness

**DOI:** 10.1101/2023.03.03.530931

**Authors:** Richard E. Pimentel, Gregory S. Sawicki, Jason R. Franz

**Author notes:** **Corresponding Author:** Jason R. Franz, Ph.D., 10206C Mary Ellen Jones Building, CB 7575, Chapel Hill, NC 27599.

## Abstract

Aging elicits numerous effects that impact both musculoskeletal structure and walking function. Tendon stiffness (k_T_) and push-off propulsive force (F_P_) both impact the metabolic cost of walking and are diminished by age, yet their interaction has not been studied. We combined experimental and computational approaches to investigate whether age-related changes in function (adopting smaller F_P_) may be adopted to mitigate the metabolic consequences arising from changes in structure (reduced k_T_). We recruited 12 young adults and asked them to walk on a force-sensing treadmill while prompting them to change F_P_ (±20% & ±40% of typical) using targeted biofeedback. In models driven by experimental data from each of those conditions, we altered the k_T_ of personalized musculoskeletal models across a physiological range (2-8% strain) and simulated individual-muscle metabolic costs for each k_T_ and F_P_ combination. We found that k_T_ and F_P_ independently affect walking metabolic cost, increasing with higher k_T_ or as participants deviated from their typical F_P_. Our results show no evidence for an interaction between k_T_ and F_P_ in younger adults walking at fixed speeds. Individual lower body muscles showed unique effects across the k_T_ and F_P_ landscape. Our simulations suggest that reducing F_P_ during walking would not mitigate the metabolic consequences of lower k_T_. Wearable devices and rehabilitative strategies can focus on either k_T_ or F_P_ to reduce age-related increases in walking metabolic cost.

**Author Summary:** Our muscles and tendons are affected by aging. Tendon stiffness and push-off forces both impact the energy cost of walking, which in turn increases with age. We investigated whether age-related changes in function (less push-off force) may be adopted to mitigate the metabolic consequences arising from structural changes (lower tendon stiffness). Reducing push-off force during walking would not mitigate the metabolic consequences of lower tendon stiffness. Wearable devices and rehabilitative strategies can focus on either tendon stiffness or push off intensity to reduce age-related increases in walking metabolic cost.

## Introduction

Older adults consume energy roughly 10-30% faster than young adults to walk at the same speed or cover the same distance.(1–4) There are a number of morphological, biomechanical, neural and biochemical factors that may contribute to these higher metabolic costs. However, a recent narrative review implicated the potential interplay between age-related changes in series elastic tendon stiffness (a structural change) and push-off intensity (a functional change) during the propulsive phase of walking.(5) Model estimates suggest that lower tendon stiffness (k_T_) yields shorter muscle fascicle lengths, decreasing the economy of force generation and increasing muscle metabolic cost.(6,7) Similarly, empirical data in younger adults show that walking with diminished push-off intensity, measured via the peak anterior/propulsive component of the ground reaction force (i.e., F_P_), also increases the metabolic cost of walking.(8) Both of these factors (i.e., decreased k_T_ and reduced F_P_) are characteristic of elderly gait and have been independently studied in the context of walking economy. However, interactions between k_T_ and F_P_ and any resultant effects on the metabolic cost of walking have yet to be explored.

Although some discrepancies exist in the comparative literature, most human studies show that older adults exhibit lower k_T_ and higher maximal strain during force-matched functional tasks compared to young adults.(9,10) Most reports focus on the Achilles tendon due to its relevance to walking metabolic cost and accessibility for *in vivo* imaging. Age-related decreases in Achilles k_T_ associate with lower walking performance (shorter 6-minute walk test distance) in older adults.(11,12) This supports the role of elastic energy storage and return as a vital mechanism to minimize the metabolic cost of walking.(13,14) Altered k_T_ has direct influence on the mechanics and economy of muscle contractions which, at least for the Achilles tendon, may influence push-off behavior. When walking at the same speed, older adults display a diminished soleus muscle operating range compared to young adults.(15) Could walking with a reduced F_P_ mitigate the metabolic penalty we pay for reduced k_T_?

Walking function, particularly that during the push-off phase, arises from the interaction between muscle activity, muscle mechanics, and tendon elastic energy storage and return. During steady-state walking at preferred speeds, these muscle-tendon dynamics are tuned to optimize movement economy. We perform significant mechanical work during push-off – predominantly via muscle-tendon units (MTU) spanning the ankle – to propel the body forward, which exacts a metabolic cost to transition from one step to the next. Reduced F_P_ among older adults increases walking metabolic cost by placing higher demand on more proximal leg muscles to perform mechanical work.(16) Specifically, demand for mechanical power normally accommodated by distal MTUs spanning the ankle is redistributed to more proximal MTUs spanning the hip.(16,17) This has metabolic consequences because, unlike those spanning the hip, MTUs spanning the ankle are uniquely designed for economical force production during walking with relatively shorter fascicles and longer tendons.

Although individuals can increase k_T_ in response to mechanical stimuli from exercise(18,19), it can be challenging to quantify the role k_T_ plays in modulating walking whole-body metabolic cost. Fortunately, musculoskeletal modeling(20–22) overcomes some of these methodological challenges. Our lab(6) and others(11,23) have used such models to reveal that decreasing k_T_ elicits shorter muscle fiber lengths, requiring higher activations and thus higher metabolic costs to meet the task demands of walking. However, prior studies have only augmented Achilles’ k_T_ (rather than *all* of the tendons in the simulated lower limb), even though there is little experimental data to suggest that age-related decreases in k_T_ are limited to tendons spanning the ankle. Furthermore, limiting altered k_T_ to only the Achilles tendon may disguise other compensatory muscle actions.

Our purpose was to quantify the individual and combined effects of k_T_ and F_P_ on walking metabolic cost in total and at the individual-muscle level. Our central motivation was to determine whether walking with reduced F_P_ mitigates the metabolic penalty of reduced k_T_. We hypothesized that: 1) decreasing k_T_ would contribute to higher metabolic costs during walking; and 2) k_T_ and F_P_ would significantly interact to affect the metabolic cost of walking. More specifically, we thought that reducing F_P_ may offer a way to mitigate higher metabolic costs anticipated with decreased k_T_. We also explored our experimental effects on muscle activation and fiber length to probe the mechanisms underlying the k_T_, F_P_, and metabolic cost landscape. Our anticipated results are intended to provide valuable insight into the tendency of older adults to walk with smaller F_P_ and diminished ankle push-off, and how clinicians, scientists, and engineers might design devices and interventions to overcome the burden of inefficient walking.

## Results

### Total Metabolic Cost

We found significant main effects of k_T_ (p<0.001, η_p_^2^= 0.423, Fig. 2 horizontal axis) and F_P_ (p=0.014, η_p_^2^= 0.244, Fig. 2 vertical axis) on total metabolic cost. We did not find a significant interaction between k_T_ and F_P_ (p=0.162, η_p_^2^=0.111). In general, total metabolic cost increased as k_T_ decreased (ε_o_ increased) or as F_P_ deviated from the Norm intensity.

**Figure 1:**
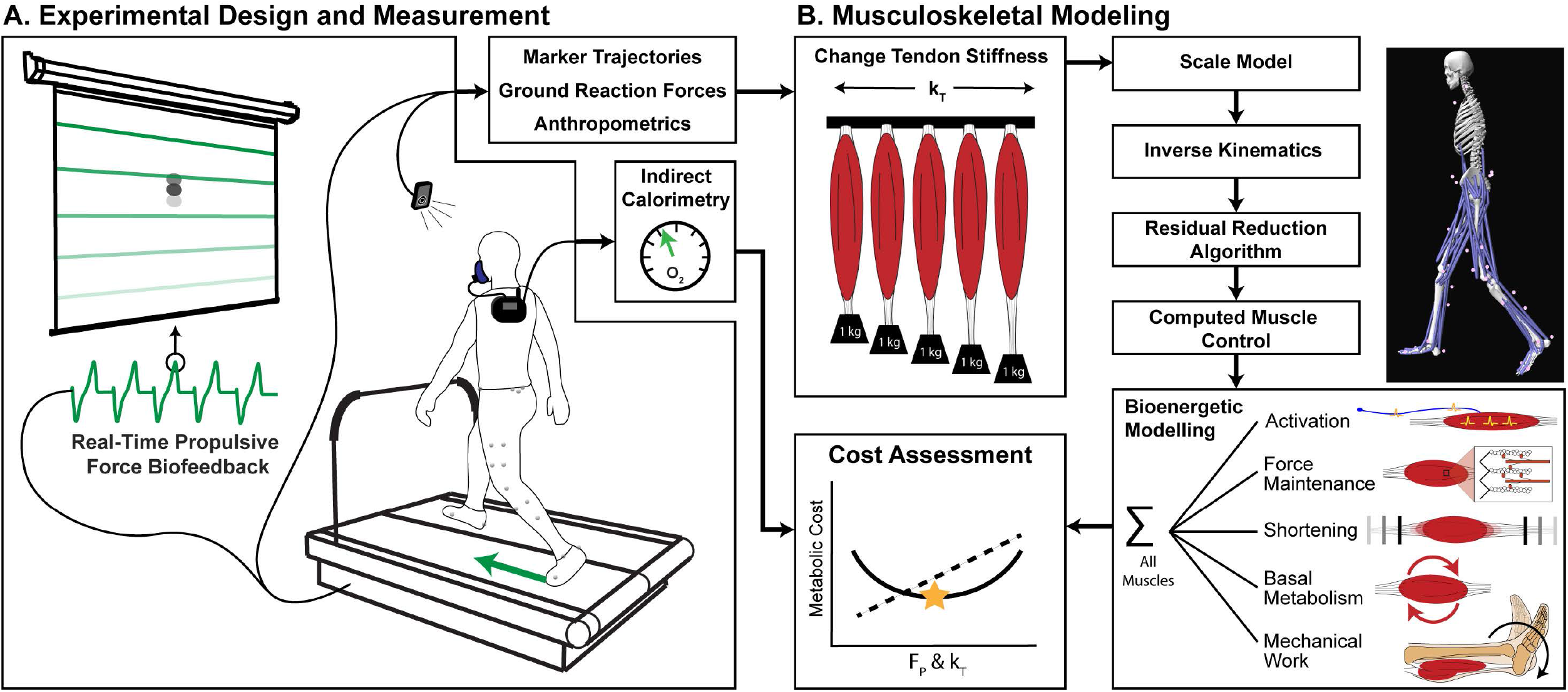
An overview of the (A) experimental and (B) computational methods to examine how tendon stiffness (k_T_) and propulsive force (F_P_) affect the metabolic cost of walking. In our experimental design (A), we asked participants to walk at their preferred speed while targeting specific F_P_ using visual biofeedback. We simulated their movement in numerous musculoskeletal models at a range of k_T_ levels (ε_o_ = 2%, 3.3% (model default), 4%, 6%, and 8%) and estimate d the metabolic cost for each condition (k_T_ & F_P_).

**Figure 2:**
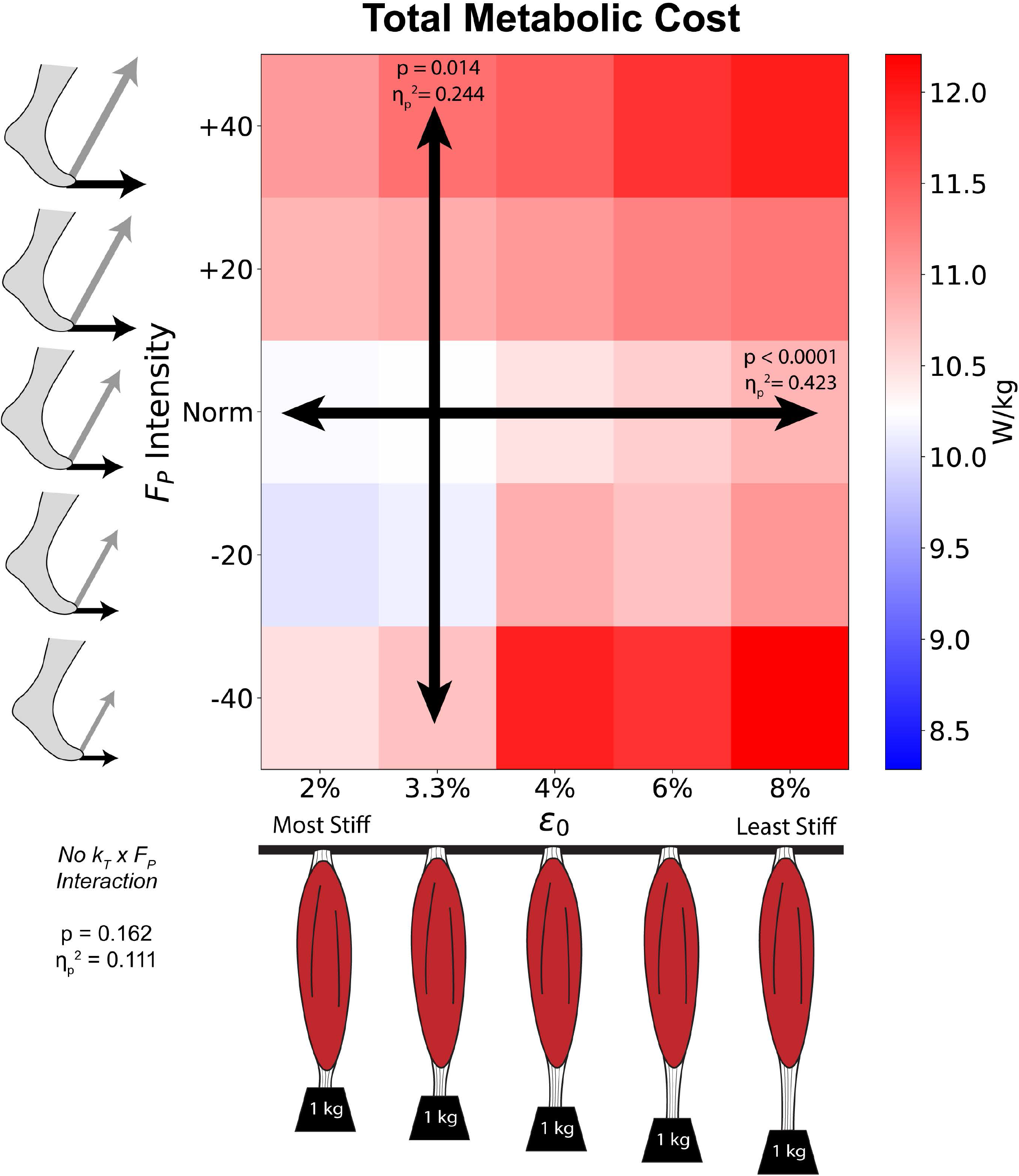
We show how average total metabolic cost varies across *k_T_* (horizontal axis) and F_P_ intensity (vertical axis). This heatmap is color coded for the reference metabolic cost (default k_T_ and Norm F_P_ intensity) to be displayed in white, with higher costs in red and lower costs in blue. We found significant ANOVA main effects separately for k_T_ (horizontal arrow) and F_P_ (vertical arrow), but no interaction (no diagonal arrow) between them.

When viewed across the gait cycle (Fig. 3), we found that k_T_ and F_P_ affect instantaneous metabolic cost differently across various phases of the gait cycle. We found significant main effects of k_T_ (ε_o_) during early stance (10-16% gait cycle), late stance (48-52% and 55-60% gait cycle) and late swing (92-100% gait cycle). Alternatively, we found significant main effects of F_P_ during mid-to-late stance (~25-32% and 40-45% gait cycle) and mid-to-terminal swing (72-80% and 97-100% gait cycle). In general, the effects of k_T_ and F_P_ on total metabolic cost occurred at different times of the gait cycle, not simultaneously.

**Figure 3:**
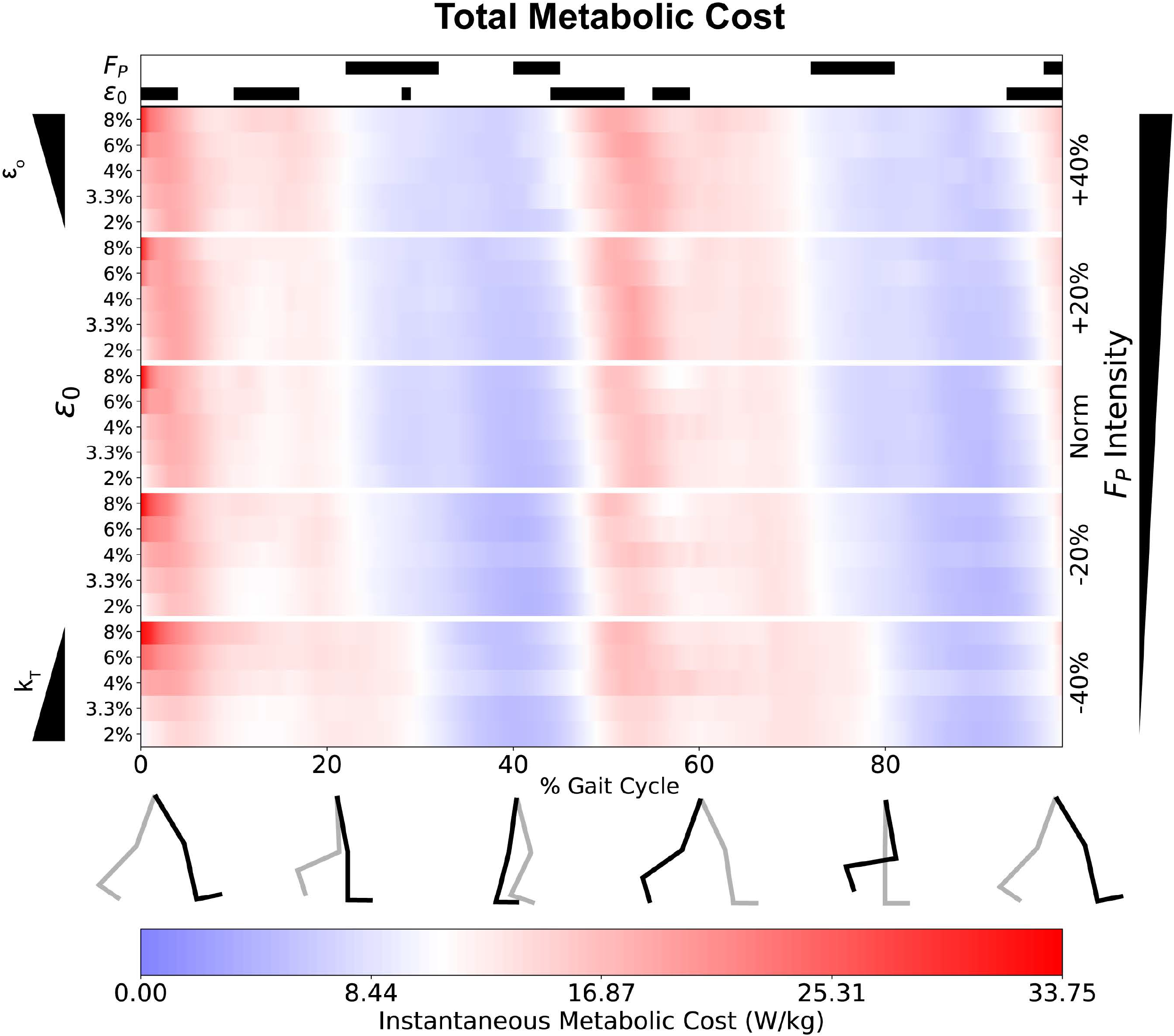
We show instantaneous, whole-body metabolic cost as a percentage of the gait cycle, across both k_T_ (left minor axes) and F_P_ (right major axis). The average metabolic cost at default k_T_ and Norm F_P_ is normalized to white. We display higher costs in red, and lower costs in blue, with the color intensity profiles even between the two. At the top of the figure, we show periods with significant main effects for k_T_ (or ε_o_) and F_P_. The black horizontal bars indicate a significant repeated measures ANOVA main effect via statistical parameter mapping.

### Individual Muscle Metabolic Costs

Table 1 shows the metabolic costs for all modeled muscles, ranked in order of energy consumption, and how their consumption varied as a function of k_T_ and F_P_. Ten different muscles contributed >4% (>2% unilateral) to total metabolic cost. The three highest energy-consuming muscles (*glut_med, rec_fem, and soleus*) each contributed >8% (>4% unilateral).

Figure 4 graphically summarizes the effects of k_T_ and F_P_ for the top 12 most costly muscles, which accounted for 70.8% (35.4% unilateral) of the total metabolic cost at default k_T_ and Norm F_P_. Eight of top 12 individual-muscle contributors to total metabolic cost showed a main effect of k_T_ (horizontal arrows), including all three muscles spanning the ankle and three out of the five muscles spanning the knee. Nine of the top 12 contributors to total metabolic cost showed a main effect of F_P_ (vertical arrows), including six of the seven muscles spanning the hip (all but *rec_fem*). Finally, five of the top 12 most energy-consuming muscles showed an interaction effect (diagonal arrows), including all three muscles spanning the ankle.

**Figure 4:**
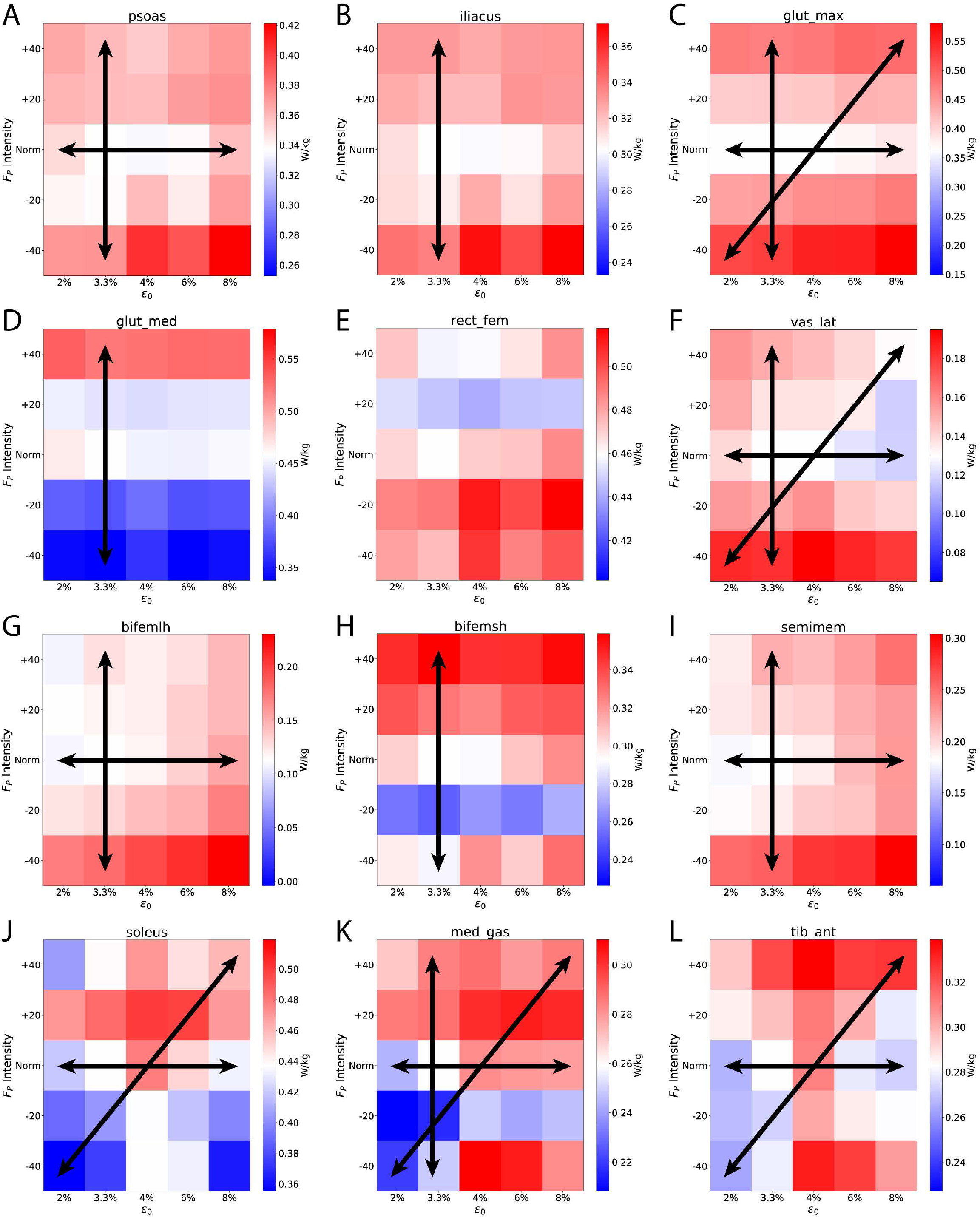
Individual muscle metabolic costs respond uniquely across k_T_ and F_P_. In this figure, we show the top 12 lower-body muscles that contribute to walking metabolic cost (Table 1). We oriented the heatmaps with proximal musculature (hip) towards the top, and distal musculature (ankle) towards the bottom. Like Figure 2, we normalized each heatmap for the default k_T_ and F_P_ values (3.3% and Norm, respectively) to be shown in white, with higher costs in red and lower costs in blue. Within each heatmap, we show significant ANOVA main effects via horizontal, vertical, and diagonal arrows indicating significant effects for k_T_, and F_P_, and interaction, respectively.

Figure 5 shows how the individual muscle metabolic costs vary across the gait cycle. Although we did not see phases of interactions between k_T_ and F_P_ for total metabolic cost, we did see such interaction effects at the individual-muscle level during specific instances of the gait cycle (gray shaded regions along the top bar). For example, on average, we found significant interactions between k_T_ and F_P_ for *soleus, glut_max, tib_ant, and med_gas* (Table 1). We can see these interactions during early-to-mid stance for *glut_max* (Fig. 5C); early stance, push-off, and late swing for *soleus* (Fig 5J); intermittently throughout stance phase for *med_gas* (Fig. 5K); and push-off for *tib_ant* (Fig. 5L).

**Figure 5:**
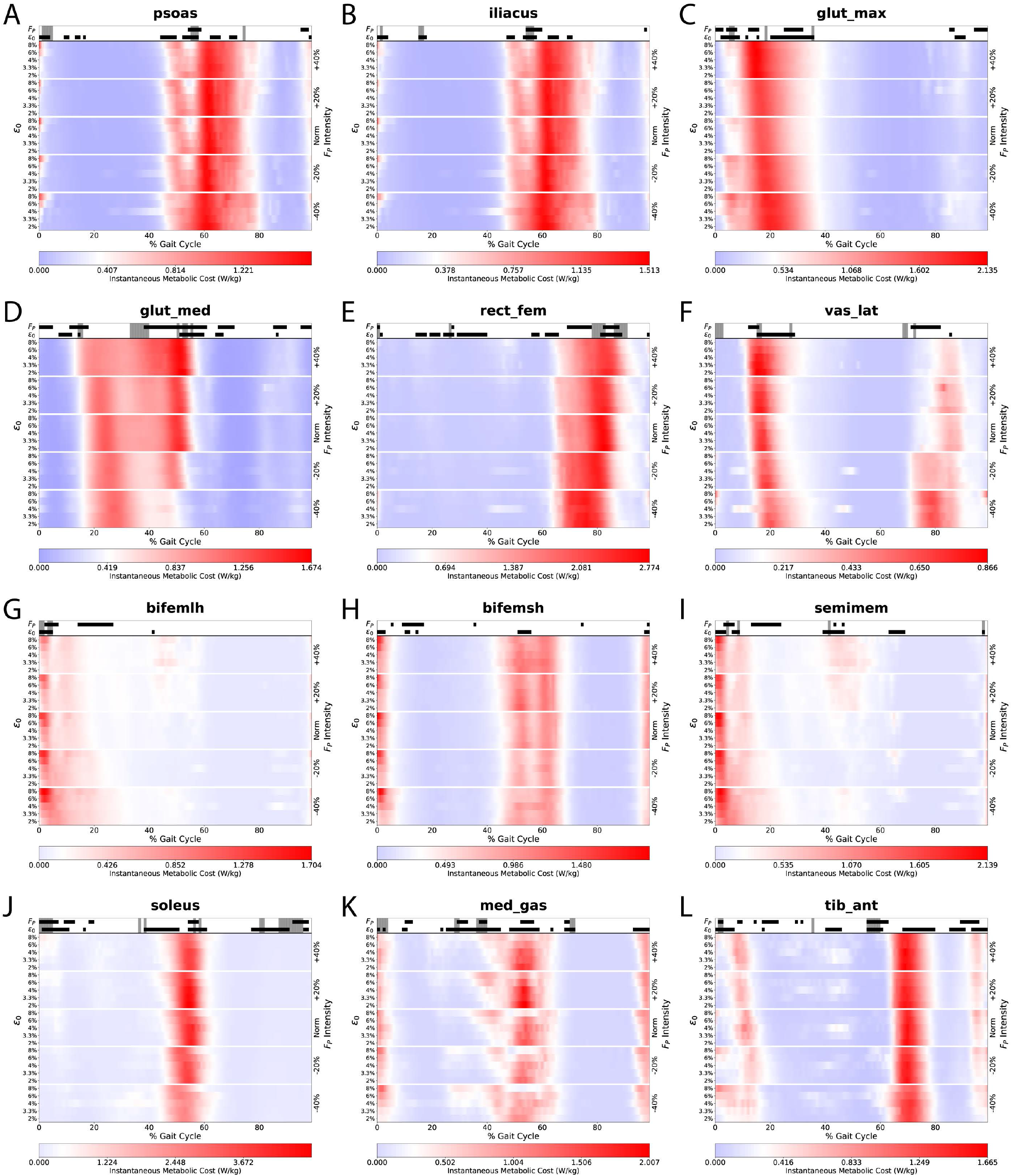
Timing and intensity of individual-muscle metabolic costs change when varying k_T_ and F_P_. In this figure, we show 12 lower-body muscles as in Figure 4, now including instantaneous metabolic cost across the gait cycle. These heatmaps are designed similar to the whole-body costs in Figure 3, with k_T_ on the left minor vertical axis, F_P_ on the right vertical major axis, and relative time (% GC) on the horizontal axis. We show significant ANOVA main effects from instantaneous statistical parametric mapping using blocks (ε_o_ and F_P_) and shaded regions (interactions) along the top bar.

### Individual Muscle Fiber Lengths and Activation Levels

We found a significant association between metabolic cost and mean activation (Fig. 6A) but not for normalized mean fiber length (Fig. 6B). F_P_ had a mixed influence on the relationship between activation and fiber length (Fig. 6C), whereas k_T_ (strain) showed a strong tendency for shorter fiber lengths and higher activations (Fig. 6D, downward & rightward shift from blue [most stiff] to pink [least stiff]).

**Figure 6:**
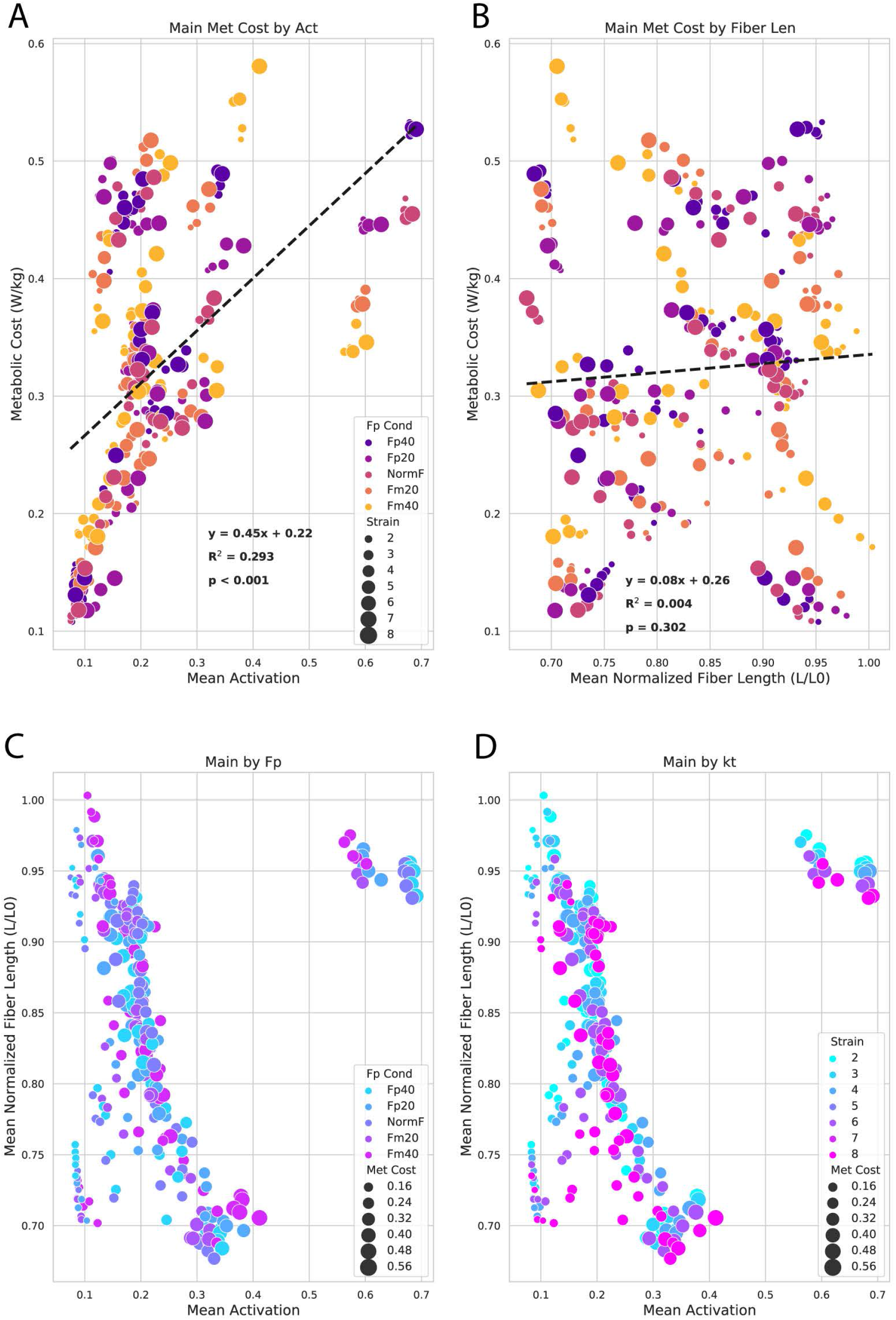
These scatterplots show general associations between the underlying muscle-tendon dynamics of activation and fiber length and their influence on the relationships between metabolic cost, F_P_, and k_T_. Each circle represents the subject-average outcome at a given activation, fiber length, metabolic cost, F_P_, and k_T_ for the top dozen individual-muscle contributors to metabolic cost (also shown in Figures 4-5). We found a significant association between metabolic cost and mean activation (A) but not for normalized mean fiber length (B). F_P_ (C) seemed to have more mixed influence on the relationship between activation and fiber length, whereas k_T_ (strain, D) showed a strong effect for shorter fiber lengths and higher activations (downward & rightward shift from blue (most stiff) to pink (least stiff)).

For additional context and transparency behind our simulations, we provide supplementary figures showing the simulated activation levels (Supplementary Figures 1 & 3) and fiber lengths (Supplementary Figures 2 & 4) for total and individual-muscle metabolic costs. We also show the average activation levels and normalized fiber lengths at default k_T_ and Norm F_P_ for these muscles in Supplementary Tables 1 & 2. On average across the gait cycle, the top dozen energy-consuming muscles (bolded muscle names) tended to have higher activation levels (higher ranks in Supplementary Table 1) and shorter fiber lengths (lower ranks in Supplementary Table 2) compared with the rest of the musculature.

## Discussion

A hybrid of experimental and computational approaches allowed us to investigate whether age-related changes in function (adopting smaller F_P_) affected the metabolic consequences arising from those in structure (having lower k_T_). This combined experiment yielded simulated instantaneous metabolic costs, providing the ability to identify muscle level responses across two of the predominant aging-related factors that contribute to increased walking metabolic cost.

Our data support our first hypothesis, that decreasing k_T_ increases the metabolic cost of walking. Conversely, based on total estimates of metabolic energy cost, we reject our second hypothesis. We found no interactions between k_T_ and F_P_ in simulations of walking at a constant speed in young adults. Specifically, k_T_ and F_P_ each independently affect walking metabolic cost, with no evidence that walking with smaller F_P_ may be adopted to mitigate the metabolic consequences arising from reduced k_T_.

By exploring individual muscle contributions to total metabolic cost, we show unique inter-muscular responses to alterations in k_T_ and F_P_. In the following sections, we further interpret these cumulative results from young adults walking across a range of F_P_ intensities in the context of characteristic changes in structure and gait function in older adults. These findings highlight the importance of structural components (k_T_) and functional behavior (F_P_) as determinants of walking economy.

### Effects of Simulated Changes in Tendon Stiffness

It is well documented that k_T_ decreases with age and physical inactivity. Lower k_T_ associates with worse walking performance(11,12) generally via two mechanisms: directly via changes in tendon elastic energy storage, return, and MTU power generation, and indirectly by compelling shorter muscle lengths. For example, altering gastrocnemius and solus reference tendon strain *in silico* from 3-11% yields vastly different tendon and muscle lengths during stance, and thus the timing and magnitudes of tendon and muscle power.(6) Meanwhile, operating a less-stiff tendon elicits shorter muscle lengths for a given force output and MTU length, thereby requiring high excitations at a metabolic penalty.(6)

Altering only the Achilles k_T_ in older and younger adults yields relatively small effects (~1.5% change from baseline) on whole-body metabolic cost across a range of walking speeds.(23) Although the Achilles tendon and triceps surae musculature may be most impacted by changing k_T_, we cannot assume that only the Achilles tendon would be affected by altered k_T_. Thus, we allowed for uniform changes in k_T_ across all MTUs in our musculoskeletal models. In doing so, we found a rather large effect size for k_T_ (η_p_^2^ = 0.423, explaining ~42% of the variance in metabolic cost).(33) When considering Norm F_P_ alone, reducing k_T_ (increasing ε_o_ up to 8%) yielded a 5% increase in metabolic cost on average (10.2 W/kg at 3.3% ε_o_ vs. 10.8 W/kg at 8% ε_o_). Our findings agree well with the literature that reducing k_T_ increases the metabolic cost of walking.(6,34)

We can see instantaneous effects of k_T_ on total metabolic cost particularly during the beginning of the push-off phase (i.e., around 50% of the gait cycle, Fig 3). At that instant, simulations with lower k_T_ (i.e., 6-8% ε_o_) showed an earlier increase in metabolic cost than those with higher k_T_ (i.e., 2-4% ε_o_). Breaking this down to individual muscles, this k_T_ effect on metabolic cost arises from ankle extensors (i.e., *soleus* and *med_gas*) and hip flexors (i.e.,*psoas* and *iliacus*). We also saw main effects of k_T_ on activation for *med_gas* (Supp. Fig. 3 K) and on fiber length for all four muscles (Supp. Fig. 4 A, B, J, K). Operating against their least stiff tendons, the *soleus* and *med_gas* showed large metabolic effects of k_T_, functioned at shorter fiber lengths, and did not lengthen much during mid-stance phase (10-50% gait cycle). Conversely, operating against their stiffest tendons, these same muscles exhibit lengthening behavior during mid stance, and functioned at longer fiber lengths during push off. Qualitatively looking at muscle dynamics, k_T_ also seemed to show a direct impact on mean activation and fiber length demonstrating a clear shift towards shorter fiber lengths and higher activations (Fig 6D). Our walking simulations and metabolic outcomes are consistent with the majority of outcomes from other studies, finding that individual muscle actions and metabolic cost during walking are highly sensitive to changes in k_T_.(18,23,35,36)

### Effects of Changing Propulsive Force

In this study, we used F_P_ as our proxy for functional changes, due to its strong association with walking speed(37,38) and hallmark decline among older adults.(39–42) We recently discovered empirical evidence that a diminished F_P_ increases metabolic cost, explained via the distal-to-proximal redistribution of muscle workload.(16) Not surprisingly, walking with larger F_P_ at a fixed speed, at least among younger adults, also increases measured walking metabolic cost. Our whole-body bioenergetic predictions support these earlier measurements.

Interestingly, we did not see an effect of F_P_ on total metabolic cost during push-off phase (50-60 % gait cycle, Fig. 3). Similar studies that increased/decreased propulsion found increases in neuromuscular drive to the ankle extensors.(43,44) In agreement with the literature, we do see individual muscle effects (for metabolic cost, activation, and fiber length) during this phase for the hip flexors (*psoas* and *ilacus*), hip abductor (*glut_med*), and ankle extensors (*soleus* and *med_gas*). These individual muscle outcomes reveal the compensatory costs of walking with altered F_P_. For example, walking with greater F_P_ exacts higher ankle extensor metabolic costs. Conversely, walking with a diminished F_P_ requires the hip flexors to compensate with higher metabolic costs to drive hip flexion. As a particularly interesting outcome, *glut_med* (a hip abductor), operated at higher costs when walking with larger F_P_ (Figure 4D & 5D). We suggest this may relate to an increased need for hip stability while transmitting larger forces from the ankle extensors to the body’s center of mass.(45)

### Relevance to Precision Rehabilitation

Our results show that k_T_ and F_P_ do not interact in their effects on the metabolic cost of walking – at least in young adults. While this is a novel finding, it is most relevant when placed in the context of walking among older adults. Specifically, our data suggest that smaller observed Fp in older adults would not be an effective mitigation strategy to conserve energy to counteract lower k_T_. A logical extension of that interpretation is that interventions designed to reduce the metabolic cost of walking in older adults are not subject to a trade-off and can independently address either or both reduced k_T_ and diminished F_P_.

We envision several viable options for solutions that could address the metabolic consequences of reduced k_T_ and diminished F_P_. For example, plyometric training may help increase k_T_(19,46) and biofeedback-based gait retraining may help improve F_P_.(47) Thus far, training strategies to reduce the metabolic cost of walking among older adults have been generally unsuccessful, primarily because they focused on increasing muscle strength, rather than k_T_.(48)

Opportunities remain to simultaneously address the metabolic consequences of both reduced k_T_ and diminished F_P_. For example, ankle exoskeletons could be designed to augment ankle joint stiffness and provide supplemental F_P_ with potential to reduce the metabolic cost of walking.(35,49) In addition, exoskeletons that can provide chronic (i.e. employable over weeks to months) wearable powered assistance or resistance on-demand could be used to interleave phases of on-line gait retraining to improve volitional Fp with phases of scheduled resistance to increase muscle strength and tendon stiffness.(50) Ultimately, this stresses that holistic design, iterative functional testing, and personalized prescriptions may be needed to implement effective wearable devices or rehabilitative therapies to combat walking inefficiency and deteriorating functional ability to maintain overall health in our aging population.

### Limitations

First, it is either very time consuming (weeks of plyometric training) or highly unethical to conduct a study to alter k_T_ in human subjects. We relied on the combination of experimental procedures and simulations to explore the metabolic cost-k_T_-F_P_ landscape. A noted by others(51), simulations cannot exactly replicate the motions and forces a participant may produce at both the specified k_T_ and F_P_. Thus, our experiments may not offer exactly the appropriate constraints for estimating changes in muscle dynamics of actual participants with altered k_T_.

Second, our discoveries are, by design, inferred from data-driven musculoskeletal simulations rather than via direct empirical measures. Our simulations reported high and potentially implausible metabolic costs at the beginning of the gait cycle, relatively early in each simulation’s run time. We suspect these outcomes are simulation artifacts rather than reliable predictions. For this reason, we opted not to discuss or interpret any findings during very early stance phase (<5% gait cycle). Additionally, like most forward dynamic simulations, our musculoskeletal dynamics and metabolic cost outcomes depend on optimization algorithms with objective functions that seek to minimize the muscle activation squared. Although these simulation methods are supported by experimental evidence(26,52), human neuromechanics may not always align with their outcomes.

Third, we designed our biofeedback paradigm to prompt changes in F_P_ while at a constant speed, which also resulted in changes in stride length.(8) We did not characterize the effects of stride length specifically in this study. However, because older adults exhibit reduced stride length(53) as well as F_P_, we contend that the outcomes reported here are relevant to our target population. We have previously quantified and discussed at length these interactions between F_P_ and stride length and their potential impact on metabolic cost.(17,47)

### Conclusion

We combined computational and experimental analyses to answer the following question with clinically important implications: “Does walking with reduced F_P_ mitigate the metabolic penalty of reduced k_T_?” In an experimental paradigm designed for young adults to emulate older adult walking via targeted F_P_ biofeedback across various simulated k_T_ levels, the answer is “no”. Walking metabolic cost is elevated both with reduced k_T_ or with any deviation in F_P_ and we observed no trade-off that could enable functional adaptations to overcome altered structural properties of the musculoskeletal system. Although these distinct factors may need to be quantified individually, they need not be addressed separately, as wearable devices and rehabilitative strategies could be designed to simultaneously address one or both these key factors driving metabolic cost.

## Methods

### Participants & Experimental Design

This study leverages previously published experimental data, and a detailed description of our experimental design and method can be found elsewhere.(8,17) We recruited a convenience sample of 12 (7 female) healthy young adults (*average ± standard deviation:* age: 23.3±3.1 years; height=1.74±0.12 m; mass=74.7±14.3 kg). After providing informed consent according to the local institutional review board, participants walked 4 passes in a hallway with timing gates spaced 30 meters apart to determine their preferred walking speed.

We recorded a 5-minute, habitual-walking trial for each participant on an instrumented dual-belt treadmill (Bertec Corp., Columbus, Ohio, USA) at their preferred overground speed (1.37±0.15 m/s). We measured the peak anterior ground reaction force (i.e., propulsive force or F_P_) from the final 2 minutes of the habitual walking trial by extracting F_P_ from stance phases using a 20-N vertical force threshold. For our F_P_ biofeedback, we displayed the average F_P_ from the previous 2 steps in real time on a screen in front of the participant (Figure 1A). Alongside the real-time F_P_, we displayed a target as a horizontal line (corresponding to the final 2-minute average F_P_ from the habitual trial). We then familiarized each participant to our biofeedback paradigm over a 3-minute exploration trial, ensuring that each participant could readily increase and decrease F_P_ on command prior to moving forward with the experimental protocol.

We used the average F_P_ from the final 2 minutes of the habitual walking trial as each participant’s typical F_P_ (Norm). For experimental trials, participants walked at their preferred speed for 5-minute trials while responding to biofeedback targets of their Norm F_P_ as well as ±20% and ±40% of Norm, presented in randomized order. In between each trial, participants rested in a seated position for at least 2 minutes.

Participants wore a portable metabolic measurement system (Cosmed K5, Rome, Italy) that recorded rates of expired oxygen and carbon dioxide to estimate energy expenditure. Our interest in this study involves the effect(s) of two specific factors on instantaneous metabolic cost at the whole-body and individual-muscle levels. Thus, we do not show the metabolic measurements measured via expired gases in this study. Instead, we refer readers to our previous publication, which benchmarked agreement between modeled and measured steady-state average metabolic cost (R^2^: 0.33).(17)

### Musculoskeletal Simulations

We performed musculoskeletal simulations in OpenSim(20–22) (version 4.1) to estimate individual muscle mechanics and metabolic energy costs. Using functional hip joint centers(24) and a static pose, we scaled all body segments of a *gait2392* model(25) for subject-specific anthropometrics in 3 dimensions. Following standard human musculoskeletal modeling techniques (Figure 1B) described previously(17), we performed computed muscle control simulations(26) across a range of tendon strain levels (i.e., ε_o_, tendon strain at maximum isometric force). Specifically, we simulated ε_o_ at 2%, 3.3% (model default), 4%, 6%, and 8% for all model tendons (92 in total). We changed tendon strain because we could not directly alter tendon stiffness in the musculoskeletal model.

During the simulations, we probed muscle metabolic costs using the Bhargava 2004 and Umberger 2010 models(27,28), which are readily available in OpenSim. For a conservative estimate, we report the average of these two bioenergetic models. All metabolic cost presented in this study result from the musculoskeletal models, not from indirect calorimetry. Furthermore, we refer to muscle names using the standard nomenclature from the *gait2392* model MTU actuators. Finally, we follow syntax from the metabolic models and report the sum of all modeled muscles as *Total* metabolic cost.

To follow common terminology, we use the term tendon stiffness (k_T_) generally throughout, rather than tendon strain (ε_o_). We acknowledge that this complicates the narrative due to the inverse relation between stiffness and strain (2% ε_o_ = most stiff, 8% ε_o_ = least stiff). We clarify these parameters in our figures by labeling both strain (ε_o_) and stiffness (k_T_) whenever possible.

### Data Reduction & Statistics

From a selection of gait cycles over the final 2 minutes of each trial, we reduced simulation input data (motions & forces) down to one gait cycle on each side by selecting the first left and right stride from the 10-second window with the most accurate biofeedback targeting during the two minutes (as performed previously(17)). We performed a total of 600 computational simulations (i.e., 12 subjects, 5 biofeedback targets, 5 ε_o_ values, 2 strides *[left & right]*). We averaged metabolic cost estimates bilaterally for each condition prior to statistical analysis. Outcome variables included total and individual-muscle metabolic costs, reported on average and as a percentage of the gait cycle. We simplified muscles with multiple lines of action (i.e., glut_med1, glut_med2, glut_med3) by summing each component for the whole muscle (i.e., glut_med).

To analyze the effects on stride-average walking metabolic cost, we performed two-way repeated measure analyses of variance (ANOVA) to test for main effects of and interactions between k_T_ and F_P_ at whole-body and individual-muscle levels (α=0.05). Alongside the ANOVA results, we also report partial eta squared (η_p_^2^) effect sizes. Similarly, to assess the effects on walking metabolic cost as a percentage of the gait cycle, we used statistical parametric mapping(29,30) to quantify main effects of k_T_ and F_P_ (α=0.001). We also performed Pearson correlations to explore associations between primary variables (metabolic cost, k_T_, and F_P_) and muscle-level determinants (i.e., activation and fiber length). We performed all statistical calculations using the Pingouin and SciPy packages.(31,32) For transparency and to support open science, we provide our simulation data and processing scripts at: https://github.com/peruvianox/kT-Fp-MetCost.

## Acknowledgments

We thank our participants for volunteering their time and acknowledge the NIH for funding this work (R01AG058615).

---

**Supplementary Figure 1:**
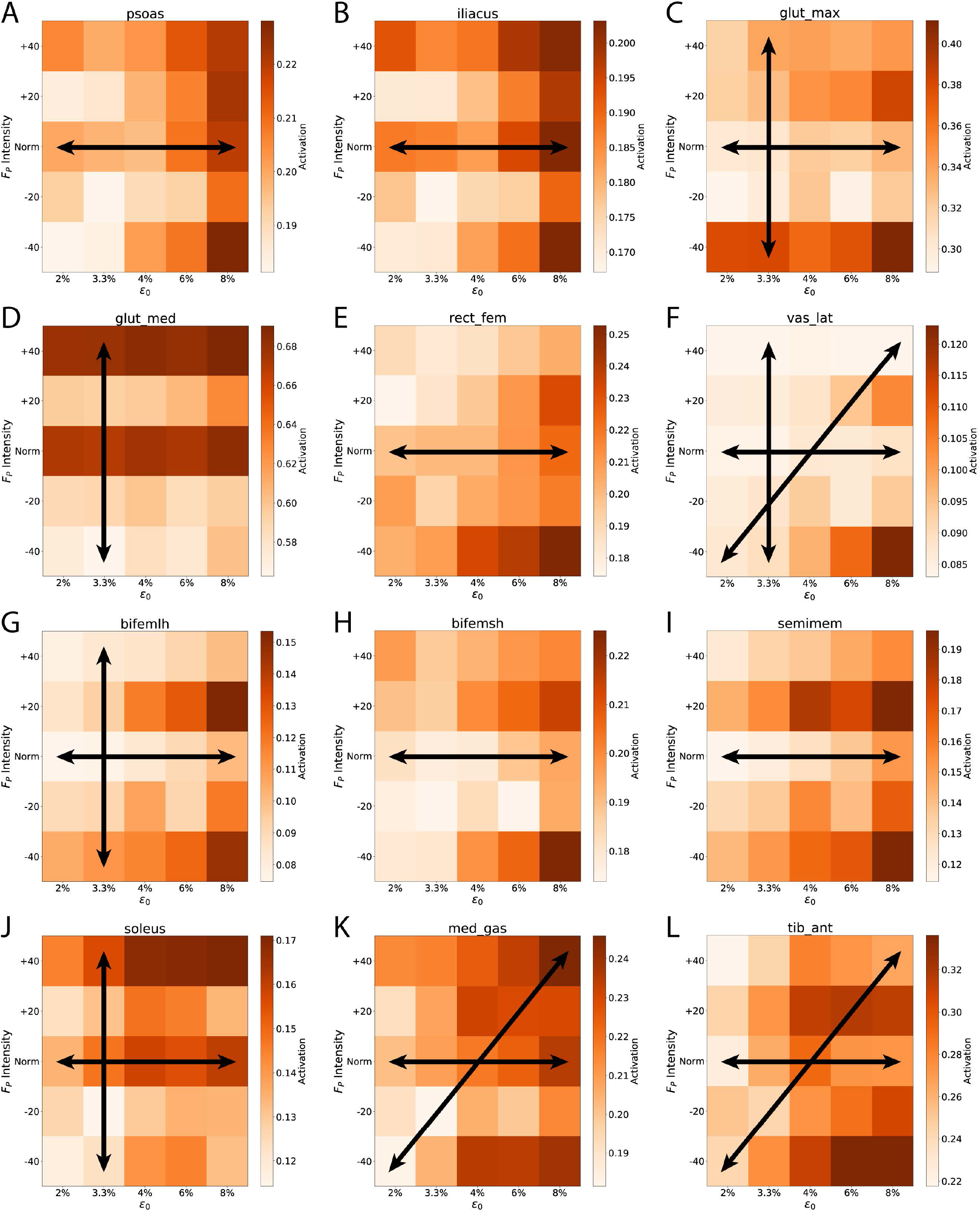
We show individual-muscle activation levels as a function of both k_T_ and F_P_. ANOVA main effects are shown via arrows (horizontal, vertical, & diagonal) similar to Figures 2 & 4. Eleven of the 12 highest energy consuming muscles showed significant effects in activation level for k_T_ (all but *glut_med*, panel D). Five of the 12 most costly muscles showed significant effects in activation level for F_P_. Three out of the 12 muscles displayed significant interactions between k_T_ and F_P_ for activation level. These data correspond with Supplementary Table 1.

**Supplementary Figure 2:**
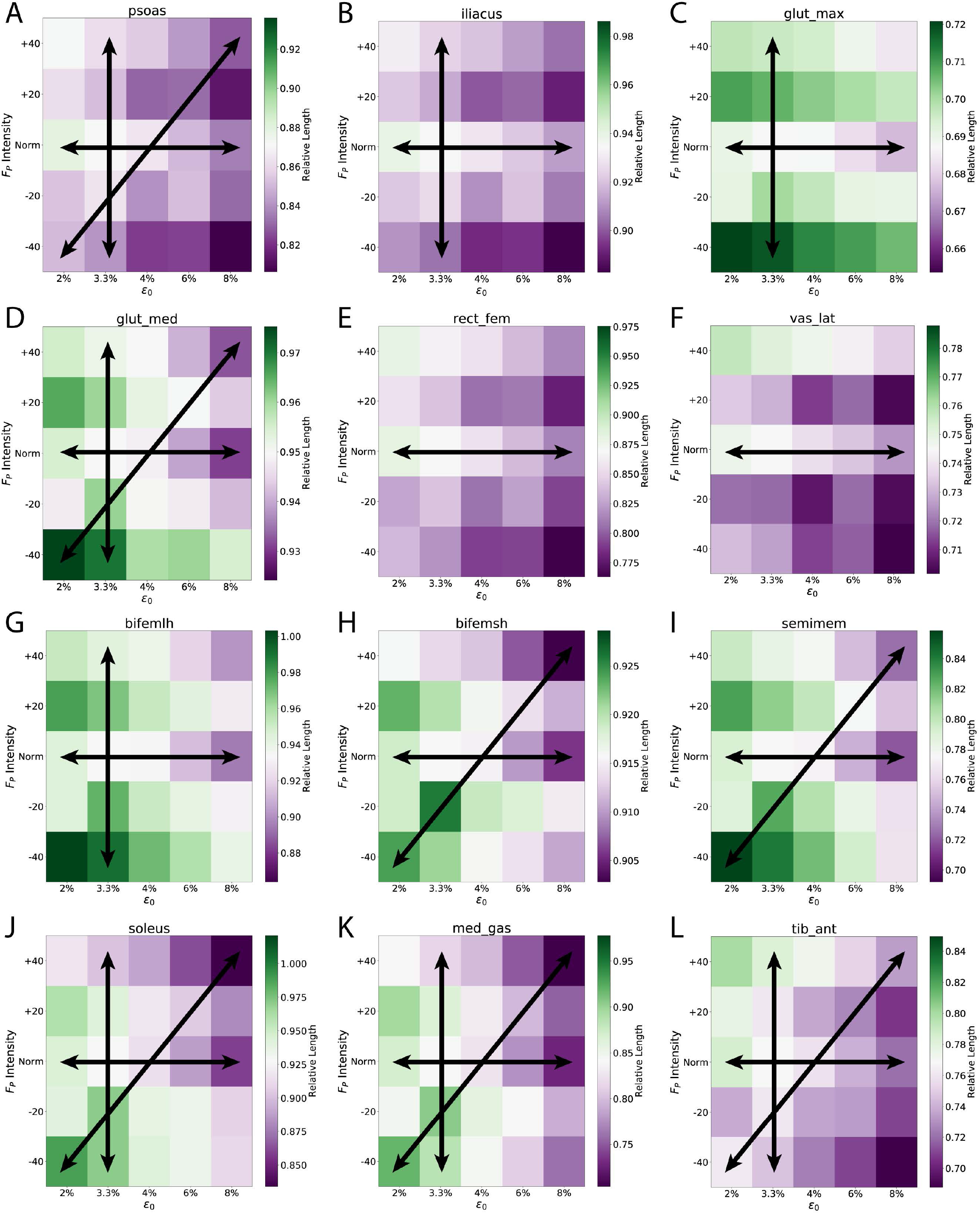
We show individual-muscle fiber lengths as a function of both k_T_ and F_P_. The fiber lengths of all 12 of the costliest muscles were significantly impacted by k_T_. Additionally, 8 of these top 12 had significant effects for F_P_, while 7/12 showed significant interaction effects. These data correspond with Supplementary Table 2.

**Supplementary Figure 3:**
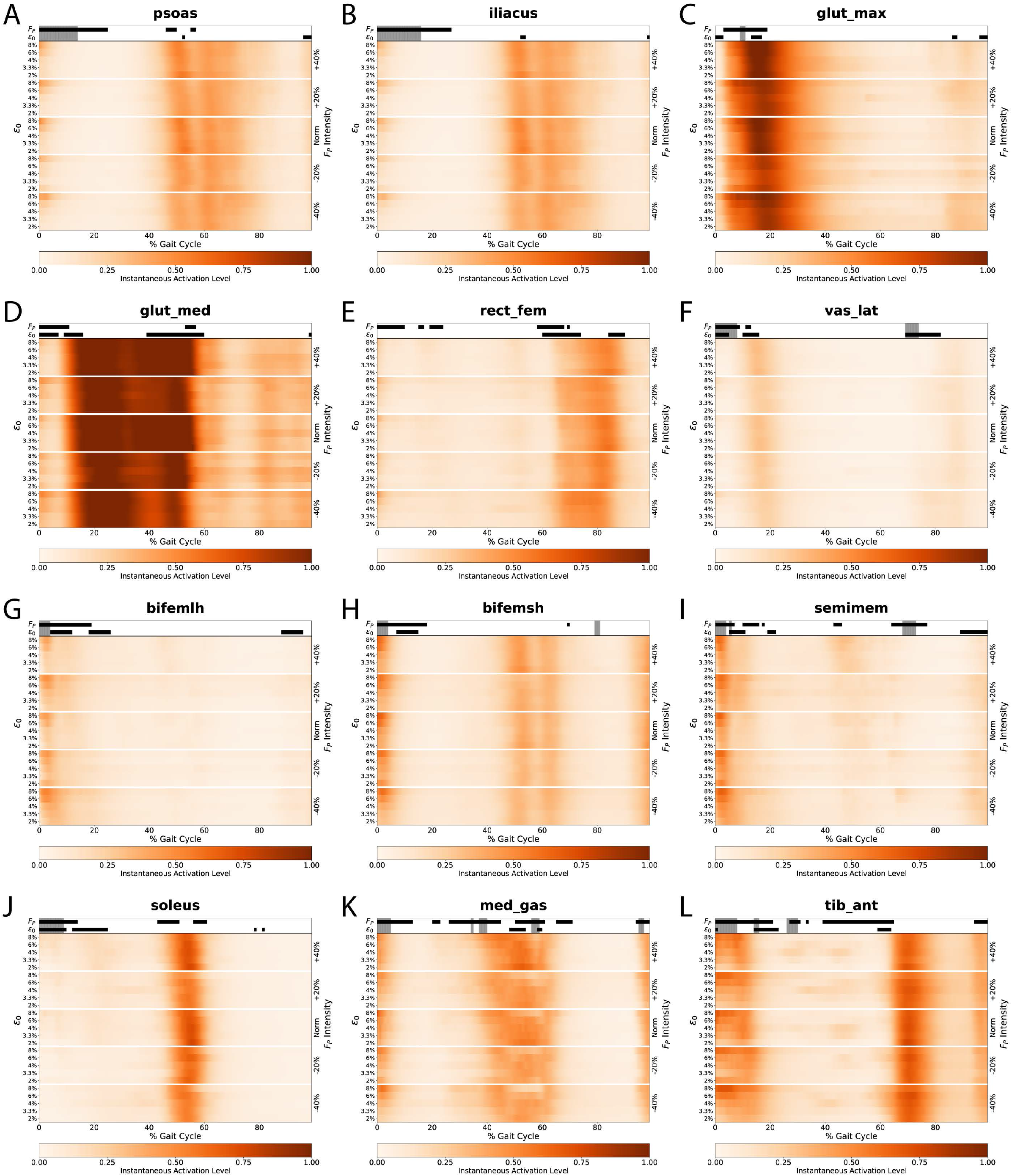
The instantaneous activation levels for highest consuming muscles highly aligned with the instantaneous metabolic costs (Figure 5).

**Supplementary Figure 4:**
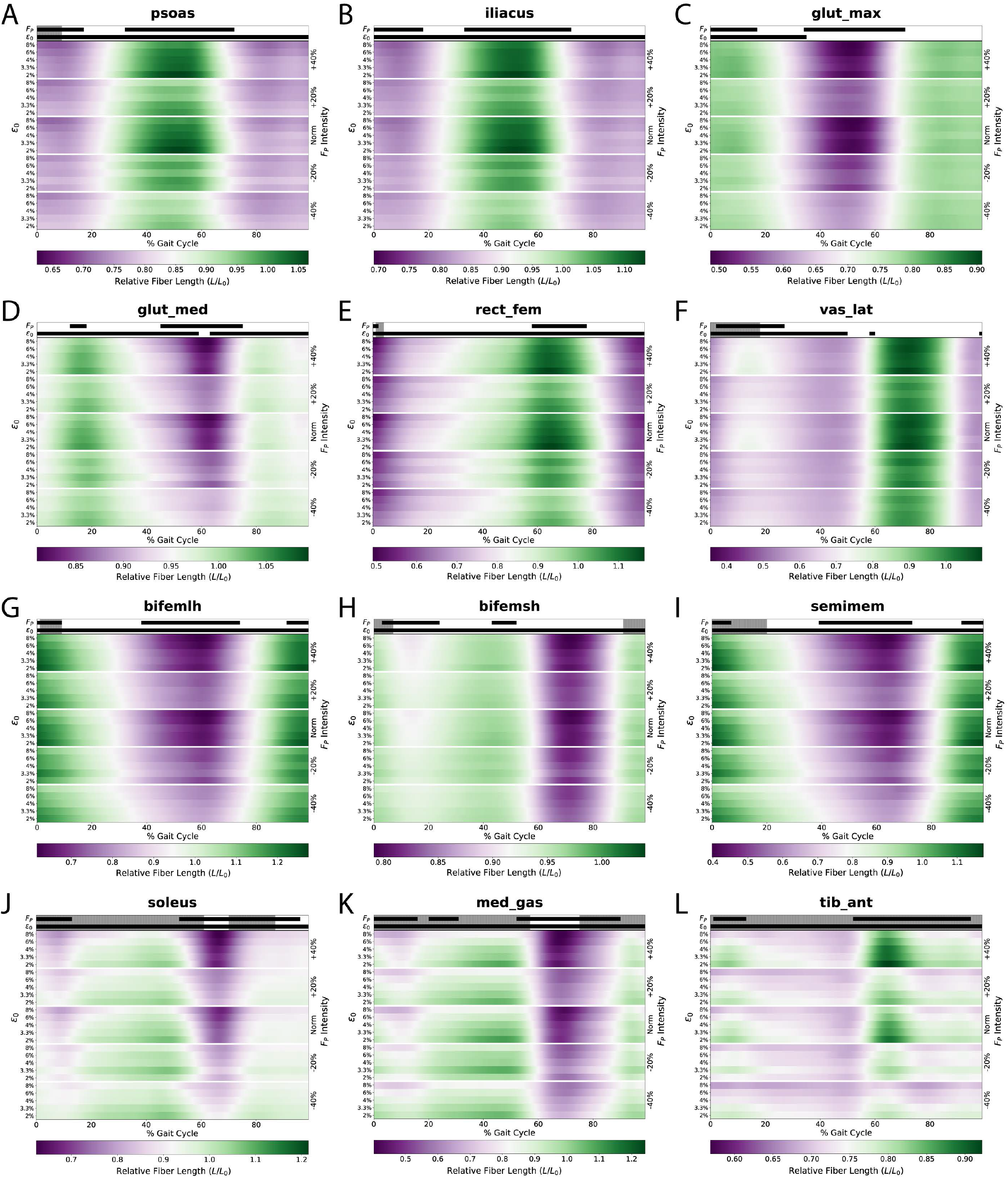
Viewing the instantaneous muscle fiber lengths, we see changes across the experimental conditions for both k_T_ and F_P_. These showcase underlying changes in muscle actions as a result from the altered k_T_ and F_P_. Large effects occur in the distal musculature, particularly for the *solues, med_gas*, and *tib_ant* (J, K, & L).

